# Molecular physiological characterization of the dynamics of persister formation in *Staphylococcus aureus*

**DOI:** 10.1101/2023.06.21.545909

**Authors:** Shiqi Liu, Yixuan Huang, Sean Jensen, Paul Laman, Gertjan Kramer, Sebastian A. J. Zaat, Stanley Brul

**Affiliations:** Department of Molecular Biology and Microbial Food Safety, Swammerdam Institute for Life Sciences, University of Amsterdam, 1098 XH Amsterdam, The Netherlands; Laboratory for Mass Spectrometry of Biomolecules, Swammerdam Institute for Life Sciences, University of Amsterdam, 1098 XH Amsterdam, The Netherlands; Department of Medical Microbiology, Centre for Infection and Immunity Amsterdam (CINIMA), Amsterdam UMC, 1105 AZ Amsterdam, The Netherlands

**Keywords:** antibiotic persistence, *Staphylococcus aureus*, proteomics, metabolomics

## Abstract

Bacteria possess the ability to enter a growth arrested state known as persistence in order to survive antibiotic exposure. Clinically, persisters are regarded as the main causative agents for chronic and recurrent infectious diseases. To combat this antibiotic-tolerant population, a better understanding of the molecular physiology of persisters is required. In this study, we collected samples at different stages of the biphasic kill curve to reveal the dynamics of the cellular molecular changes that occur in the process of persister formation. After exposure to antibiotics with different modes of action, namely vancomycin and enrofloxacin, similar persister levels were obtained. Both shared and distinct stress responses were enriched for the respective persister populations. However, the dynamics of the presence of proteins linked to the persister phenotype throughout the biphasic kill curve and the molecular profiles in a stable persistent population did show large differences depending on the antibiotic used. This suggests that persisters at the molecular level are highly stress specific, emphasizing the importance of characterizing persisters generated under different stress conditions. Additionally, although generated persisters exhibited cross-tolerance toward tested antibiotics, combined therapies were demonstrated to be a promising approach to reduce persister levels. In conclusion, this investigation sheds light on the stress-specific nature of persisters, highlighting the necessity of tailored treatment approaches and the potential of combined therapy.

**Importance:** By monitoring proteome and metabolites during *Staphylococcus aureus* persister formation under vancomycin and enrofloxacin exposure, we revealed the dynamic information of the molecular physiology of persister formation upon exposure to two different antibiotics with different modes of action. The data shows that cells that phenotypically are similarly classified as persisters, do have several molecular characteristics in common but, remarkably so, differ substantially in a significant number of other aspects of their molecular makeup. These contrasts provided valuable insights into persister eradication, which holds considerable clinical relevance.

## Introduction

Antibiotic persistence (hereinafter referred to as persistence), a non-inheritable antibiotic-tolerant growth-arrested phenotype, has been confirmed as the major cause of recurrent and chronic infections, as well as one of the main contributors to the development antibiotic resistance. Persisters were firstly discovered in *Staphylococcu aureus* populations as early as in 1942 [1] and are now found in almost all tested bacterial species [2]. When exposed to antimicrobials, cells can switch a sub population into persisters that can easily resuscitate back to normal cells with culturability and virulence after stress removal, causing recurrent infection during treatment gaps. Thus, a better understanding of the molecular physiology of persisters is expected to contribute to solving these issues.

However, despite the severe impact of persisters and the value of knowing the mechanisms of their formation, this phenotype is still puzzling. For decades, the complexity and contradictory information on persisters have been discussed in terms of their high phenotypic heterogeneity [2 – 4], the variety of mechanisms leading to their generation [5–7] and sometimes the lack of a uniform definition of persisters [8]. Regardless of the these challenges, experimentally, a biphasic kill curve is a widely agreed hallmark of persistence [9]. During a high concentration antibiotic treatment, the death of most susceptible cells results in a rapid drop in the curve regarded as phase I, followed by the survival of non-dividing persisters, indicated as phase II, the survivor plateau.

In the present work, *S. aureus*, one of the most widespread Gram-positive infectious agents and most frequent pathogen causing chronic disease, was used. By sampling at different time points of the biphasic kill curve, we studied the molecular characteristics of vancomycin or enrofloxacin treated persistent population at the protein and metabolite level, by proteome and metabolome analysis, respectively. Vancomycin is a glycopeptide antibiotic that is clinically used as a primary antibiotic of choice against gram-positive pathogens, including *S. aureus*. It inhibits the elongation and cross-linking of peptidoglycan, causing a perturbed cell wall and eventually cell death [10]. Enrofloxacin, on the other hand, is a third generation fluoroquinolone antibiotic that inhibits DNA synthesis via targeting type II-DNA gyrase and type IV-topoisomerase [11]. It is the first fluoroquinolone of choice in veterinary medicine for numerous diseases including *S. aureus* mastitis [12].

Under vancomycin or enrofloxacin exposure, our tested *S. aureus* cultures displayed a similar biphasic kill curve. In the stable persistent population, proteomics data show several common alterations in carbon metabolism, cell wall biosynthesis and the expression of ribosomal proteins. However, dynamic analysis revealed that there were major differences between the two antibiotic exposed groups. Moreover, specific molecular pathways were also found to be related to vancomycin or enrofloxacin treatment. In summary, our study highlights the generic molecular physiological makeup of persisters in *S. aureus* that were triggered by two types of antibiotic stresses. At the same time, our findings demonstrate that stress responses are highly antibiotic-specific, and not all persisters exhibit identical molecular and physiological features. These results have important implications for developing treatments for persisters, as targeting the specifically upregulated processes by an antibiotic may be pursued in require complementarying approaches with other antibiotics that can act synergistically in inhibiting persister formation.

## Results

To identify the proper sampling timepoints representing the progress of persister formation, a time-kill kinetics assay was conducted under the exposure of vancomycin or enrofloxacin (figure 1A). Biphasic killing curves showed two phases: a quick kill (phase I) that lasted for 3-4 hours and the subsequent plateau (phase II) indicating the persistent population. Within the biphasic kill curve, 24-hour treated samples were considered as stable persistent populations (hereinafter referred to as V-24h and E-24h). Vancomycin and enrofloxacin caused similar kill curves and no significant difference was found between the survival fractions in V-24h and E-24h. The proportion of persisters in V-24h and E-24h was detected through flow cytometry (figure 1B). And around 75 % of the detected events represented persisters (unpublished data), suggesting that persisters are the major population in the 24h antibiotic treated samples.

**Figure 1.**
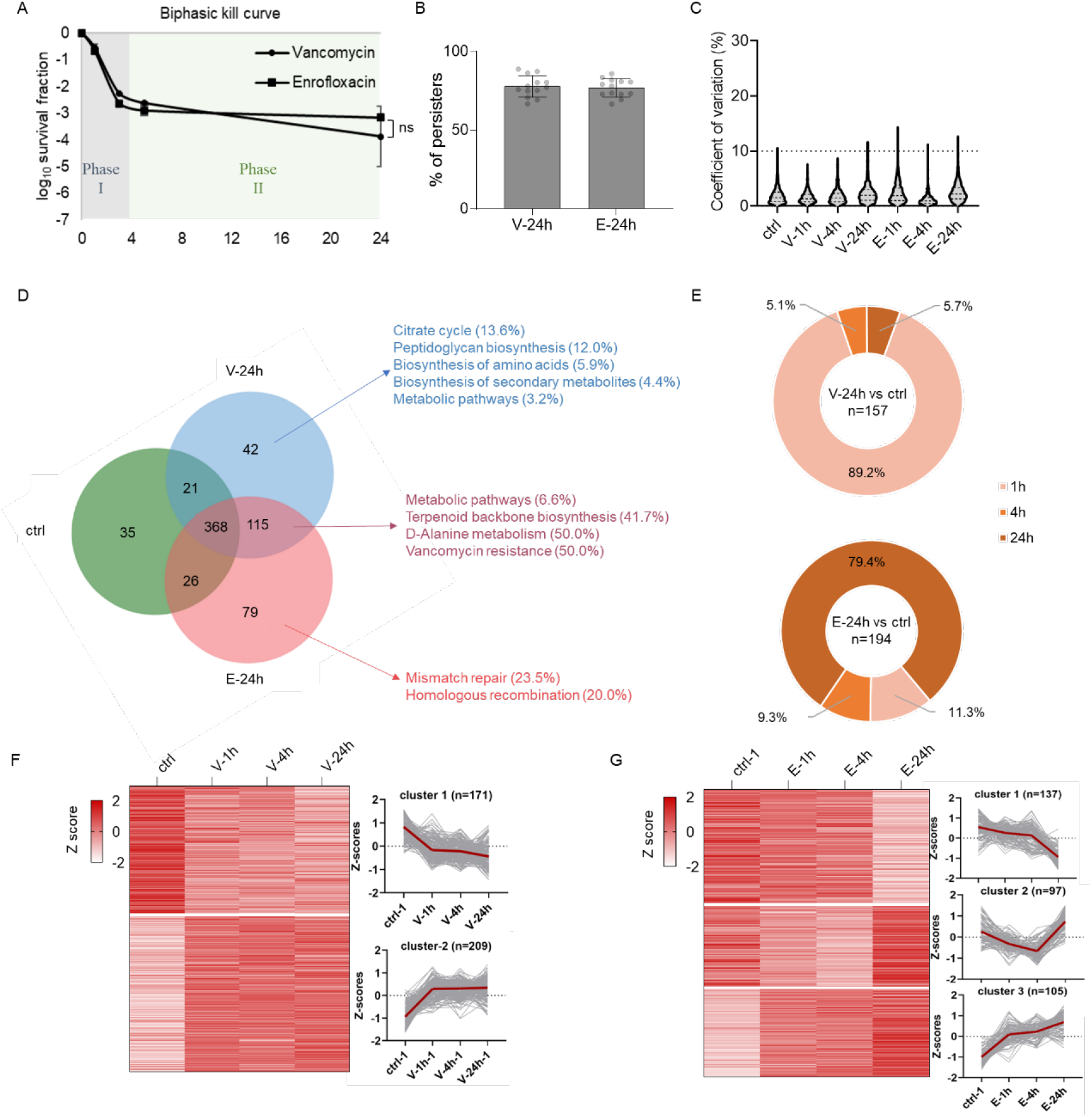
Sample characterization and unique proteome in persistent populations. **A**. Time-kill kinetics assay results in biphasic kill curves, demonstrating persister formation in a *Staphylococcus aureus* population during 24h of antibiotic exposure. Specifically, in phase I both antibiotics quickly killed around 99.9% of cells after 3-hours, leaving behind unsusceptible persisters after phase II. **B**. CFDA and PI double staining and subsequent flow cytometry analysis show that in 24h treated samples the majority of the cells (∼75 %) are persisters (our unpublished observations). **C**. OD15 cultures before and after 1h-, 4h- and 24h- antibiotic exposure were sampled for proteome analysis. The coefficients of variation between biological triplicates indicates high quantitative reproducibility. **D**. Venn diagram of identified proteins from ctrl (green), V-24h (blue) and E-24h (red), as well as enriched KEGG pathways. **E**. The present proportion of newly synthesized proteins in V-24h and E-24h compared with the untreated control at each time point. For the expression dynamics among proteins that were constitutively expressed, hierarchical cluster analysis was performed with vancomycin treated samples (**F**) and enrofloxacin treated samples (**G**). The average values are shown for each protein.

### Proteome profile dynamics during the generation of persistence

For proteomics, OD15 cultures were sampled at 1h, 4h and 24h of antibiotic exposure, representing the beginning of the fast kill process, and the beginning and the end of persistent population formation, respectively. OD15 of an early log phase culture was used as control. In total, 21 samples being 7 treatment groups with 3 replicates of each group, were harvested. Overall, 765 identified proteins present in at least two replicates in at least one group were analyzed for a general quality assessment. The average CV (coefficient of variation) of each treatment was below 3% (figure 1C), demonstrating a high quantitative reproducibility between biological triplicates. We identified 546 proteins in V-24h and 588 proteins in E-24h. Compared with 450 proteins in control cells, the Venn diagram (figure 1D) shows that vancomycin and enrofloxacin exposure triggered 157 and 194 newly synthesized proteins, respectively, among which 115 proteins were common in both V-24h and E-24h samples.

After 24h vancomycin exposure, 42 expressed proteins specific for this treatment were enriched and categorized in five metabolic pathways: citrate cycle, peptidoglycan biosynthesis, biosynthesis of amino acid, biosynthesis of secondary metabolites and metabolic pathways. Of these, dihydrolipoyl dehydrogenase (PdhD), citrate synthase (Cs) and fumarase (FumC) involved in the TCA cycle were detected and are discussed in the next section. Peptidoglycan biosynthesis suggests a protective stress response towards this cell wall targeting antibiotic. Among 115 proteins shared by the vancomycin and enrofloxacin exposed cells, three pathways that are involved in cell wall biosynthesis were enriched, suggesting that cell wall alteration can be a general stress response against antibiotics despite a different primary mode of action of antibiotics. Among 72 unique differentially expressed proteins in E-24h, two DNA repair related KEGG pathways, mismatch repair and homologous recombination, were enriched. Both of these pathways have been found essential for bacteria to survive under quinolone-induced stress conditions to combat accumulated DNA damage [13–15].

Aside from the mentioned KEGG pathways, we also observed the distinctive expression of a RelA_SpoT domain-containing protein (Q2G2L7) in V-24h, which likely leads to the generation of more ppGpp. This molecule triggers the stringent response, known to contribute to the formation of antibiotic persisters [16]. Moreover, six ATP-binding cassette (ABC) transporters were identified in E-24h. ABC transporters are related to transmembrane transport, potentially involved in the efflux of antibiotics. The expression of genes encoding ABC-transporters has also been found to be upregulated in intracellular *S. aureus* persisters surviving inside human macrophages. [17].

To understand the dynamics of the unique proteome composition under the two different stress conditions, we timed the emergence of the newly synthesized proteins during the biphasic kill curve (figure 1E). Interestingly, under vancomycin treatment, 89.2 % of these proteins were observed within the first hour of exposure and only 5.7 % after 4 hours of exposure, showing a quick response upon vancomycin induced stress perception. In contrast, in enrofloxacin-treated cultures, 11.3 % of proteins were present at the beginning of antimicrobial exposure but almost 80% of the induced proteins were only expressed after 4 hours of exposure. Similarly, hierarchical cluster analysis classified two and three clusters with distinct expression patterns from vancomycin treated samples (figure 1F) and enrofloxacin treated samples (figure 1G), respectively. Overall, these findings suggest that the proteome composition is altered differently in the response to the tested stress conditions, where vancomycin tends to cause quick proteome changes within 1h of treatment, but where most changes in enrofloxacin treated samples happen after 4h of exposure.

Next, we analyzed 368 commonly expressed proteins in control, V-24h and E-24h. Compared to untreated control samples, vancomycin induced 44 proteins and repressed 44 proteins (figure 2A). In comparison, enrofloxacin caused the overexpression of 65 proteins and the repression of 49 proteins (figure 2B). The top ten predominant proteins in each category were labelled and listed in table 1, colored in red and green, respectively. In V-24h, highly overexpressed proteins are mainly related to chaperone function (ClpB, GroEL, DnaK and DnaJ), cell wall biosynthesis (VraR, MurA) and cell membrane regulation functions (branched-chain-amino-acid aminotransferase (BCAT), acetyl-CoA acyltransferase). In E-24h, most induced proteins engage in DNA repair (UvrABC system protein A, DNA polymerase III, polydeoxyribonucleotide synthase, rNDP, dCMP kinase) and chaperone function (ClpB, trigger factor). In contrast, most listed decreased proteins in both treated groups were translation related and included ribosomal proteins. KEGG enrichment analysis of differentially expressed proteins revealed five pathways in V-24 (figure 2C): ribosome, metabolic pathway, carbon metabolism, biosynthesis of antibiotics and fatty acid metabolism. In E-24h (figure 2D), altered pathways from 114 proteins are related to translation, carbon metabolism, biosynthesis of secondary metabolites, and DNA repair involved pathways nucleotide excision repair and mismatch repair.

**Table 1.**
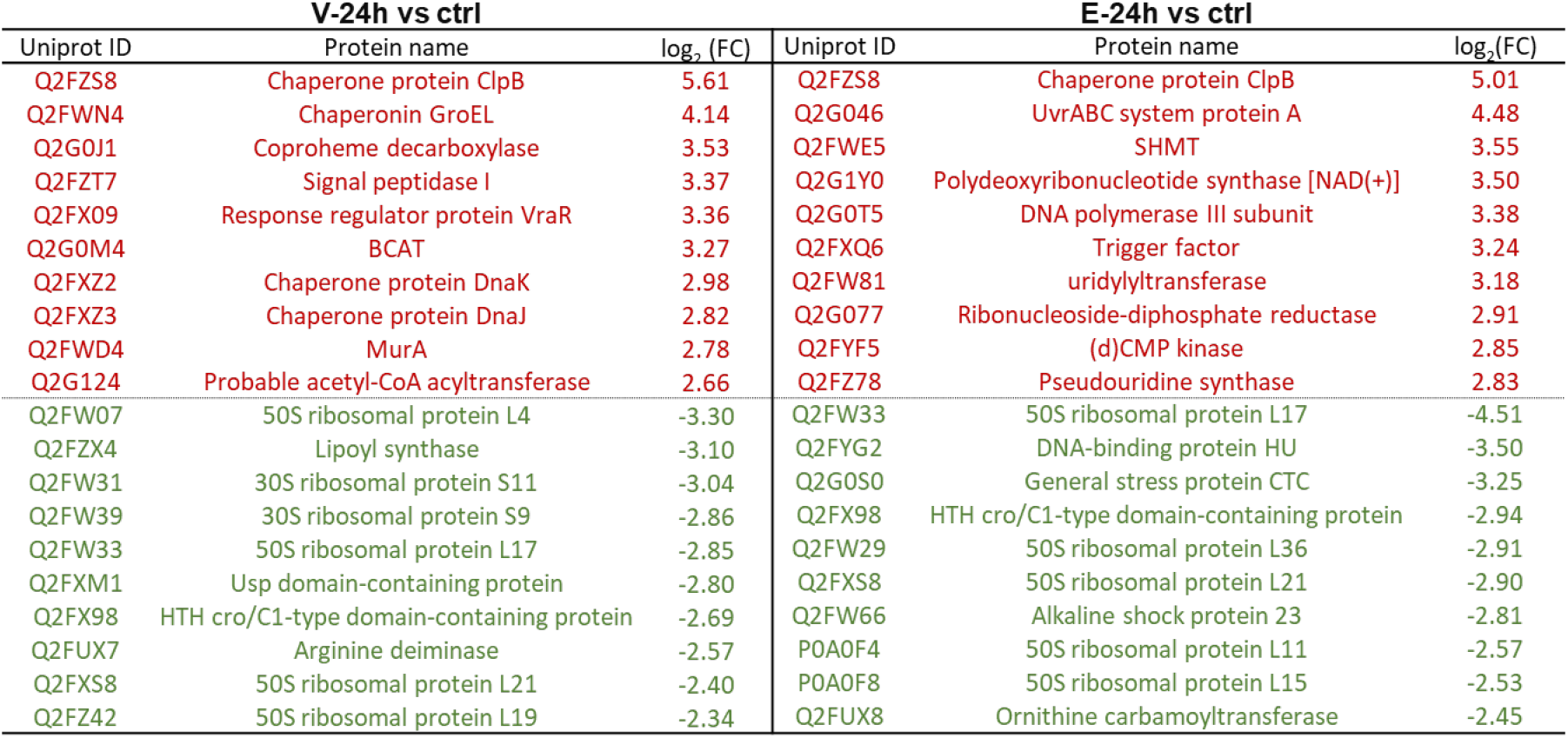
Top 10 increased proteins (red) and decreased proteins (green) in V-24h and E-24h samples compared with control samples.

| V-24h vs ctrl |  |  | E-24h vs ctrl |  |  |
| --- | --- | --- | --- | --- | --- |
| Uniprot ID | Protein name | log<sub>2</sub> (FC) | Uniprot ID | Protein name | log<sub>2</sub> (FC) |
| Q2FZS8 | Chaperone protein ClpB | 5.61 | Q2FZS8 | Chaperone protein ClpB | 5.01 |
| Q2FWN4 | Chaperonin GroEL | 4.14 | Q2G046 | UvrABC system protein A | 4.48 |
| Q2G0J1 | Coproheme decarboxylase | 3.53 | Q2FWE5 | SHMT | 3.55 |
| Q2FZT7 | Signal peptidase I | 3.37 | Q2G1Y0 | Polydeoxyribonucleotide synthase [NAD(+)] | 3.50 |
| Q2FX09 | Response regulator protein VraR | 3.36 | Q2G0T5 | DNA polymerase III subunit | 3.38 |
| Q2G0M4 | BCAT | 3.27 | Q2FXQ6 | Trigger factor | 3.24 |
| Q2FXZ2 | Chaperone protein DnaK | 2.98 | Q2FW81 | uridylyltransferase | 3.18 |
| Q2FXZ3 | Chaperone protein DnaJ | 2.82 | Q2G077 | Ribonucleoside-diphosphate reductase | 2.91 |
| Q2FWD4 | MurA | 2.78 | Q2FYF5 | (d)CMP kinase | 2.85 |
| Q2G124 | Probable acetyl-CoA acyltransferase | 2.66 | Q2FZ78 | Pseudouridine synthase | 2.83 |
| Q2FW07 | 50S ribosomal protein L4 | -3.30 | Q2FW33 | 50S ribosomal protein L17 | -4.51 |
| Q2FZX4 | Lipoyl synthase | -3.10 | Q2FYG2 | DNA-binding protein HU | -3.50 |
| Q2FW31 | 30S ribosomal protein S11 | -3.04 | Q2G0S0 | General stress protein CTC | -3.25 |
| Q2FW39 | 30S ribosomal protein S9 | -2.86 | Q2FX98 | HTH cro/C1-type domain-containing protein | -2.94 |
| Q2FW33 | 50S ribosomal protein L17 | -2.85 | Q2FW29 | 50S ribosomal protein L36 | -2.91 |
| Q2FXM1 | Usp domain-containing protein | -2.80 | Q2FXS8 | 50S ribosomal protein L21 | -2.90 |
| Q2FX98 | HTH cro/C1-type domain-containing protein | -2.69 | Q2FW66 | Alkaline shock protein 23 | -2.81 |
| Q2FUX7 | Arginine deiminase | -2.57 | P0A0F4 | 50S ribosomal protein L11 | -2.57 |
| Q2FXS8 | 50S ribosomal protein L21 | -2.40 | P0A0F8 | 50S ribosomal protein L15 | -2.53 |
| Q2FZ42 | 50S ribosomal protein L19 | -2.34 | Q2FUX8 | Ornithine carbamoyltransferase | -2.45 |

**Figure 2.**
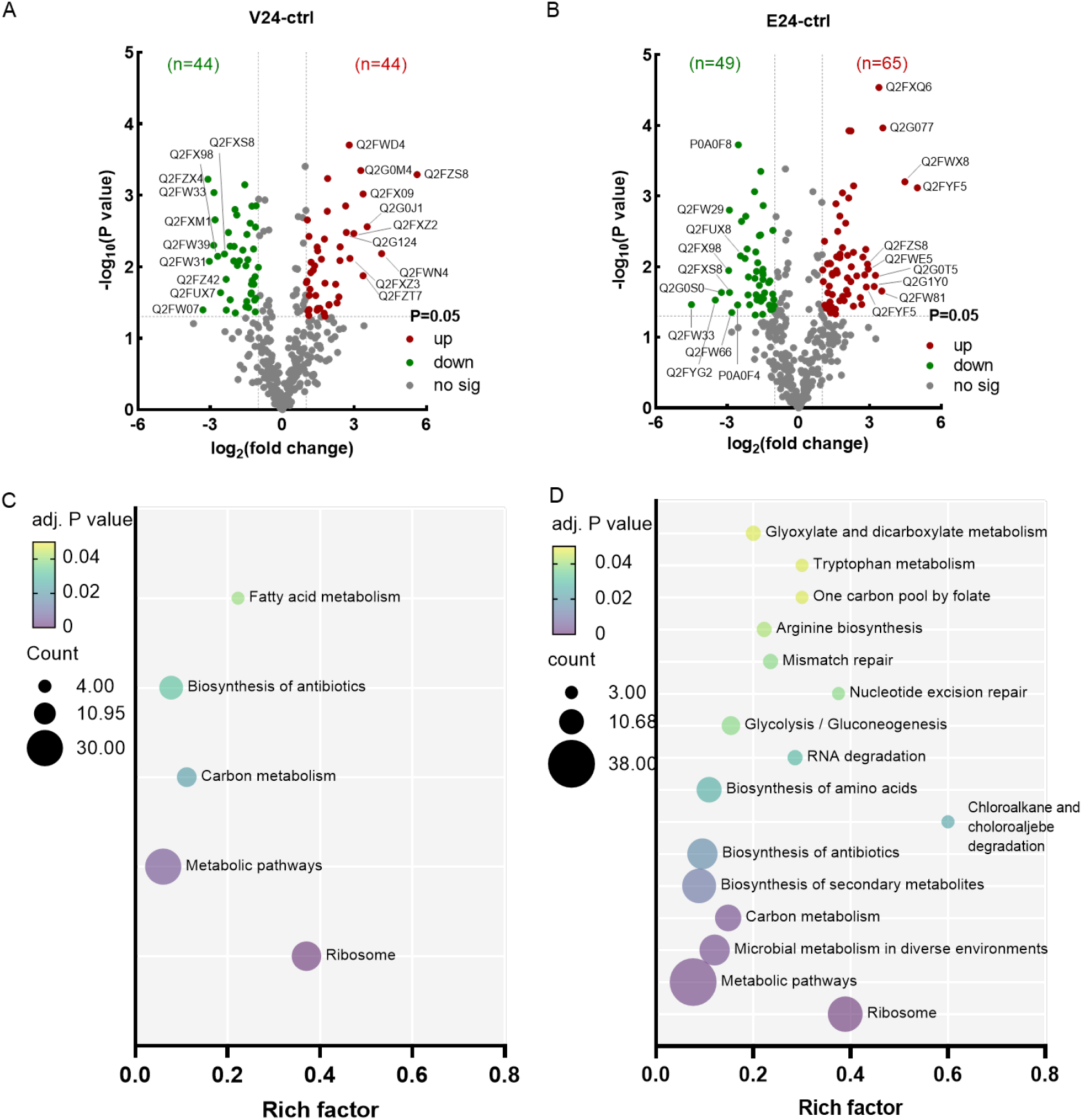
Comparative analysis between persistent populations and control cells. Volcano plots showing differentially expressed proteins (p<0.05) in the vancomycin triggered persistent populations V-24h (**A**) and the enrofloxacin triggered persistent population E-24h (**B**) compared with untreated control samples. Proteins with higher or lower expression level were labelled as red or green respectively. Uniprot ID of top-10 predominant proteins were labeled in each group. Proteins without significant differences in expression were shown as gray dots. KEGG pathway enrichment analysis was conducted with all differentially expressed proteins in V-24h (**C**) and E-24h (**D**). The enrichment factor is calculated as “identified protein numbers annotated in this pathway term”/”all protein numbers annotated in this pathway term”.

Overall, the listed proteome composition illustrates that the alteration in chaperone, carbon metabolism, cell wall biosynthesis, as well as ribosomes are common features during the formation and maintenance of persisters under both vancomycin and enrofloxacin treatment. Moreover, DNA repair is strongly associated with persistence under enrofloxacin exposure. To gain further insights into these pathways, we analyzed their dynamics and possible roles in the generation and maintenance of persisters.

### Central carbon metabolism

Central carbon metabolism is essential in both energy production and the provision of precursors for other essential cellular processes. Interesting pathways with expression level of relevant enzymes in V-24h and E24h were visualized in figure 3A. The dynamics of stated enzymes are visualized in a heatmap (figure 3B) shown in red and green for higher and lower expression compared with untreated samples, respectively. In general, in V-24h and E-24h, glycolysis was differently regulated. Meanwhile, the expression of most citrate cycle (TCA cycle) enzymes was reduced but pyruvate oxidation, pentose phosphate pathway (PPP), and one-carbon metabolism (OCM) were induced under both stress conditions. The details are discussed in the following paragraphs.

**Figure 3.**
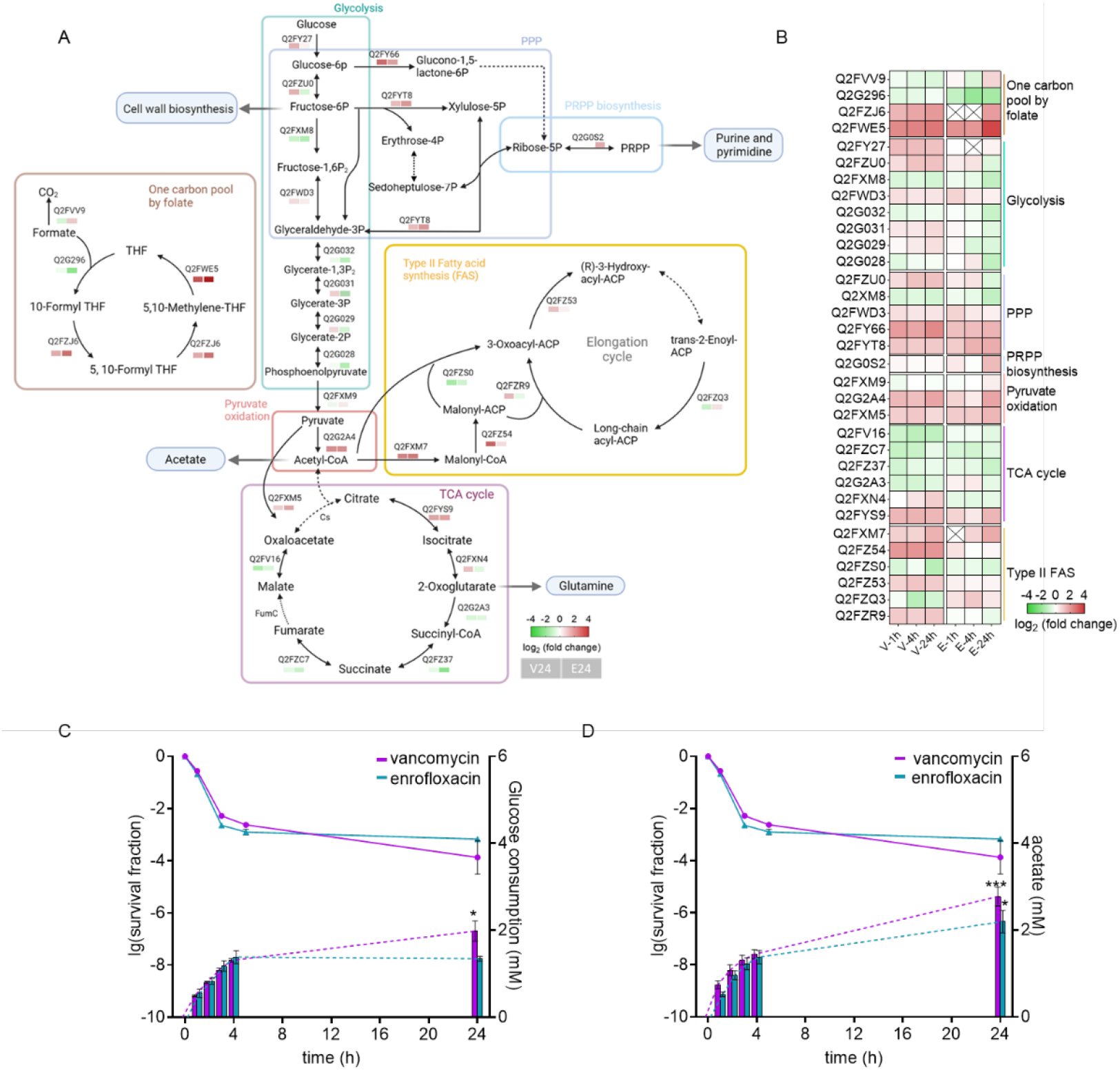
Alteration in central carbon metabolism during vancomycin and enrofloxacin exposure. **A**. Schematic map showing the fold change of relevant enzymes (labeled with UniProt protein ID) in both V-24h and E-24h. PPP: pentose phosphate pathway. **B**. The dynamics of involved proteins during the biphasic kill curve. In both **A** and **B**, green filled squares mean lower expression level while red ones are increased compared with the control group. A cross means not detected. To detect the carbon metabolic activity during antimicrobial exposure, HPLC was conducted to measure the glucose consumption (**C**) and acetate excretion (**D**). Biphasic kill curves from figure 1A are shown together with HPLC dynamics in each diagram. n=6. Unpaired T-test was used to detect the difference between 4h- and 24h-treated samples in each treatment group. *: p<0.05. ***, p<0.001.

Glycolysis is the initial pathway of utilizing glucose, the primary carbon source in culture medium. Proteomics data shows that glycolysis-involved proteins were overexpressed since the first hour of vancomycin treatment but were gradually repressed in cells exposed to enrofloxacin, indicating a difference in carbon source consumption. To investigate this, we conducted HPLC analysis to monitor glucose consumption (figure 3C) and acetate production (figure 3D) during antimicrobial exposure. Together with the biphasic kill curves (referenced from figure 1A) in each diagram, it is clear that during the initial 4 hours where most of the killing occurred, surviving cells were still able to quickly consume glucose and secret acetate in both conditions with similar activity. After this timepoint, glucose consumption ceased in enrofloxacin persisters and less amount of acetate was generated. Meanwhile, however, vancomycin persisters still had active, though relatively slower, carbon metabolic reactions.

Glycolysis is also an integral precursor provider in *S. aureus* to generate macromolecules such as amino acids, lipids and cell wall precursors [18]. For example, enzyme activities in V-24h lead to possible accumulation of fructose-6-phosphate, a glycolytic intermediate for cell wall biosynthesis [19]. Pyruvate is another important precursor generated from glycolysis to form acetate, oxaloacetate (OXA) and acetyl-CoA. Acetyl-CoA carboxylase carboxyltransferase (Q2FXM7) was elevated in both stress conditions to initiate type II fatty acid biosynthesis (FAS II). Interestingly, in V-24h, the lower expression of FabI (Q2FZQ3) which reduces trans-2-acyl-ACP to saturated long chain acyl-ACP, suggests that more unsaturated fatty acids were synthesized. During enrofloxacin treatment, FabF (Q2FZR9), catalyzing the first condensation step of the elongation cycle, gradually decreased. This suggests that the synthesis of long chain fatty acids was repressed in cells that became enrofloxacin persistent.

Under laboratory conditions and exponential growth, TCA cycle enzymes are often repressed or dispensable for viability [20]. Here in the growth arrested persistent populations, most enzymes of the TCA cycle were, as expected, repressed. Still, several enriched enzymes can be seen in the treated groups, which might be due to their role in the provision of precursors. For instance, vancomycin triggered the expression of citrate synthase (Cs), the initial enzyme of the TCA cycle, causing possible 2-oxoglutarate (2-OG) accumulation. Consistently with the literature [21], this vancomycin triggered 2-OG buildup can contribute to the steady generation of glutamine which eventually benefits cell wall biosynthesis. Vancomycin also uniquely induced the expression of FumC that was also overexpressed in persistent *S. aureus in vivo*, leading to fumarate level decrease and, by that, glycolysis activation [22].

PPP was elevated in both V-24h and E-24h, which is able to replenish NADPH that was consumed by the indicated pathways such as branched-chain amino acid biosynthesis, fatty acid biosynthesis, glutamine biosynthesis, as well as one carbon (1C) metabolism. Additionally, PPP also links to phosphoribosyl pyrophosphate (PRPP) biosynthesis, which is the rate limiting step in purine and pyrimidine biosynthesis [23]. In E-24h, enhanced synthesis of PRPP could aid in *de novo* nucleotide synthesis which is strongly linking to DNA replication, survival and antibiotic persistence in *S. aureus* [24,25]. Last but not least, folate based OCM was induced in both stress conditions, and tetrahydrofolate (THF) accumulation was expected based on the enzyme expression pattern. THF was found to directly participate in crucial metabolic activities including the synthesis of nucleotide, purine and amino acids [26]. Besides, in *E. coli*, the overexpression of 5-formyltetrahydrofolate cyclo-ligase (Q2FZJ6 in *S. aureus*) promotes persister formation upon ampicillin and ofloxacin exposure [27].

In summary, this described metabolic rewiring suggests that persisters, despite being growth arrested, alter their metabolic pathways to favor energy needs and the provision of precursors for essential cellular processes in response to antimicrobial treatment. Additionally, this adaption may also allow persisters to conserve energy and resources until conditions become favorable for regrowth.

### Chaperones

Chaperone proteins play a critical role in facilitating the proper folding of proteins in all living cells [28]. Their importance in persister formation and resuscitation was proven in several bacterial species including *S. aureus* [29–32]. In our study, we identified ten chaperone proteins that were overexpressed in almost all treated samples (figure 4A), with ClpB being the top overexpressed protein in both vancomycin-and enrofloxacin-persistent populations (see table 1). The ClpB-DnaK system has been shown to be vital for persister survival and resuscitation due to its ability to clear protein aggregates [31]. We also found increased ClpP and ClpX that associate with ATPase ClpC to form ClpPC and ClpPX proteases and facilitate persister formation via the degradation of antitoxins [33]. ClpX is also essential for cell viability in sub-optimal environments including the presence of oxidative stress [34,35], which may explain the steady increase of ClpX under enrofloxacin exposure. In addition to the Clp proteases, other chaperones such as DnaK, DnaJ, and GroEL were also overexpressed and are crucial in bacterial protein folding upon exposure to antibiotic stress [36,37]. The deletion of DnaK leads to perturbed stress response and decreased persistence [38]. In contrast, ClpL, cis/trans isomerases trigger factor (TF) and PrsA are with little known function or evidence in terms of antibiotic persistence. But TF was reported to promote biofilm formation and interact with ClpB [39] while PrsA was also involved in VraSR-regulated cell wall stress response [40].

**Figure 4.**
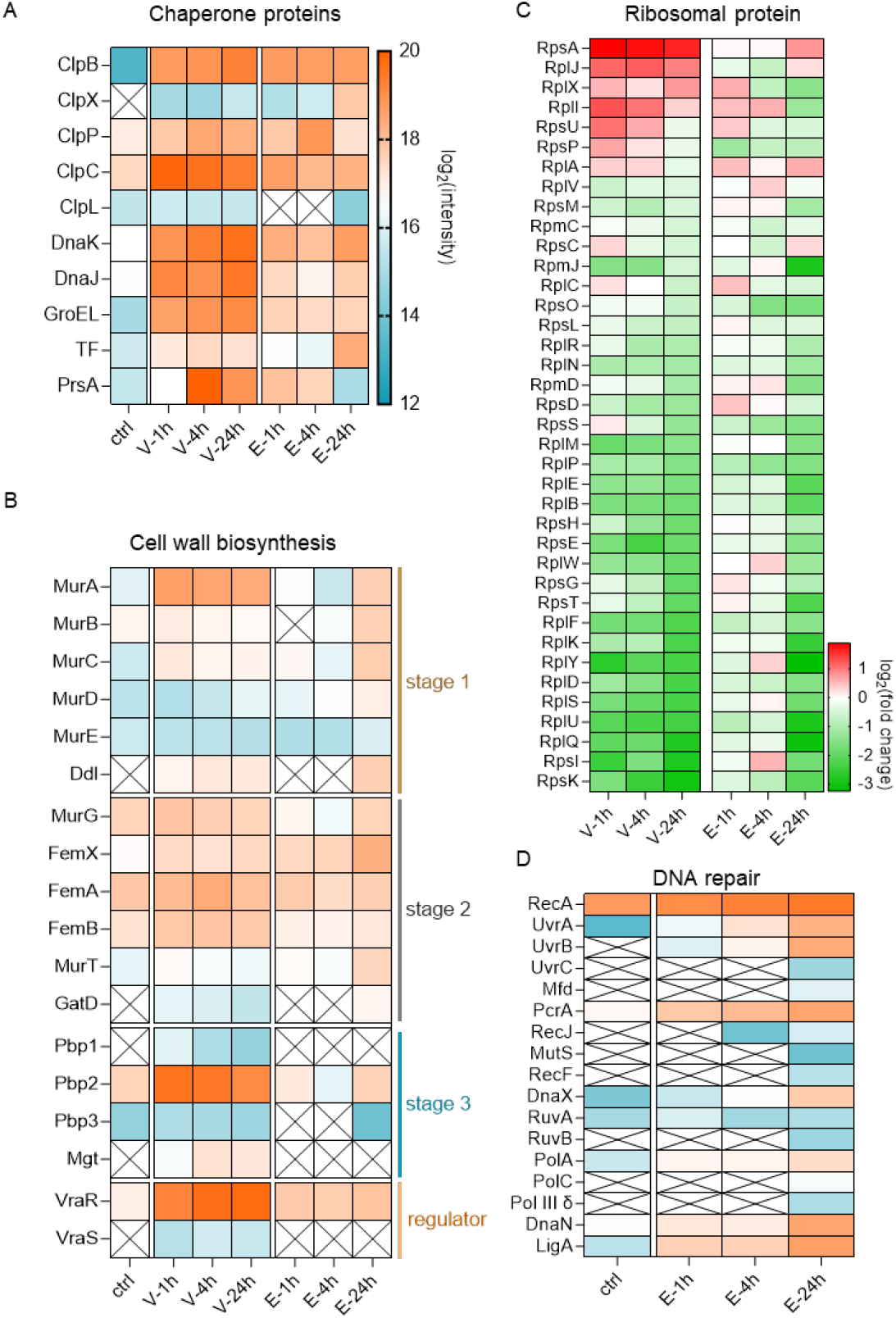
Dynamics of the abundance of (**A**) chaperone protein, (**B**) cell wall biosynthesis related proteins, (**C**) ribosomal proteins and (**D**) DNA repair related proteins. For **A, B** and **D**, the expression level of each protein are shown in color ranging from blue (low expression) to orange (high expression). A cross means not detected. In (**C**), the expression levels of ribosomal proteins are shown as log_2_(fold change) compared with control, in which green represents decreased proteins and red shows increased proteins.

Additionally, compared with untreated control samples, the increase in chaperone levels were seen within the first hour of vancomycin and enrofloxacin exposure, indicating the essential function of chaperones under the given stress conditions. Besides, vancomycin stimulated slightly more chaperones than enrofloxacin, which suggests a difference between these two persistent populations. These assumptions were supported by isolating and visualizing intracellular insoluble proteins (figure S1) during the biphasic kill curve, where aggregated proteins were observed within the first hour of incubation in all treated samples. Interestingly, the aggregation levels within vancomycin treated samples increased along the incubation while the enrofloxacin-treated samples displayed a decrease in protein aggregation over time.

### Cell wall biosynthesis

In total, nineteen proteins involved in cell wall biosynthesis were identified (figure 4B). Particularly, 17 proteins participate in the three stages of biosynthesis of peptidoglycan (PG) [41,42]. The first stage involves the synthesis of the peptidoglycan precursor uridine diphosphate (UDP)-MurNAc-pentapeptide in the cytoplasm, with enzymes such as MurA-E and Ddl being detected in our experiments. In stage 2, we identified MurG, the FemXAB family, and the MurT/GatD enzyme-complex, which stimulate the assembly and membrane translocation of lipid II. In stage 3 where the cross-linking of peptidoglycan occurs, enzymes Pbp1-3 and Mgt were detected. Additionally, we found the presence of the VraS/VraR two-component regulatory system. Upon cell wall stress, VraR regulates more than 40 proteins including Pbp2 and Mgt [21], with VraS functioning as a rapid stress sensor [43].

While KEGG enrichment analysis suggests that cell wall synthesis contributes to persistence under both vancomycin and enrofloxacin exposure, different dominant mechanisms in each group are shown. During vancomycin exposure, the highly induced proteins (>5-fold compared to control cells) participate in all three stages of PG synthesis and the VraSR regulon, supporting the idea that a general enhanced cell wall biosynthesis is induced to combat vancomycin, rather than just the synthesis stage 3 where vancomycin targets. In addition, all changes happened within the first hour of vancomycin exposure, implying an instantaneously enhanced induction of a stress protective mechanism against this cell wall targeting antibiotic. In comparison, the various proteins in enrofloxacin-triggered persistent population mostly feature in stage 1 of PG biosynthesis and displayed a relatively mild but consistent increase during the treatment. Here, the lack of induction of the VraS regulon might indicate that the induced peptidoglycan biosynthesis by enrofloxacin is not due to cell wall stress.

### Ribosomal proteins

A ribosome-dependent translation block is considered as one of the hallmarks of persistence [44]. In total, we identified thirty-eight ribosomal proteins in both V-24h and E-24h. The binary logarithm of fold change compared with control was calculated and shown in the heatmap (figure 4C). In general, the amounts of ribosomal proteins were expectedly reduced. Interestingly, most proteins with reduced abundance, especially in enrofloxacin treated samples, were gradually decreased during the biphasic kill curve instead of quickly reaching their final levels. Besides, four slightly higher expressed ribosomal proteins can be seen respectively in V-24h (ribosome initiation protein RpsA, RplJ contributing to the binding of elongation factor, assembly initiator protein RplX and RplI that influences tRNA stability) and in E-24h (RpsA, RplJ, RplA that involved in releasing tRNAs, and RpsC which is essential for mRNA entrance) [45,46]. Overall, these alterations benefit persister formation by repressing unnecessary translation, but also allow cells to synthesize stress-response-related proteins and to maintain the potential for resuscitation.

### DNA repair

We identified 21 proteins related to DNA repair in E-24h (figure 4D), including SOS response inducer RecA; nucleotide excision repair related proteins UvrABC, Mfd and PcrA; mismatch repair involved RecJ and MurS, homologous recombinant proteins RecF, DnaX, RuvA and RuvB, DNA replication related proteins Pol I (PolA), Pol III subunits PolC, Pol III δ, and DnaN, as well as DNA ligase LigA to join breaks and fill the gap generated during DNA repair and replication [47–50]. Of these, SOS response is regarded as a key response inducing persister formation via repair of DNA damage, altering DNA expression and sometimes modulating the toxin-antitoxin system [51]. Upon DNA damage, the positive regulator RecA recognizes the damage and likely binds to ssDNA (single-stranded DNA) to initiate the SOS response [52]. UvrABC, RuvAB and RecF are also associated with the SOS response and are regulated by RecA. All 21 detected proteins either steadily increased during the treatment under enrofloxacin or were detected only after a few hours of stress incubation, showing their significance in both persister formation and maintenance. The different dynamics and appearance time of enzymes suggests their putative specific function in certain stages of persister formation.

### Comparison between persistent populations generated from different stress conditions

The stated comparison between treated and untreated samples suggests that the detailed protein compositions for vancomycin or enrofloxacin persistent populations are distinct. Therefore, we comparatively analyzed the proteome and metabolomic features of V-24h and E-24h.

Firstly, volcano plots (figure 5A) show that in total 107 proteins out of 483 shared proteins are differentially expressed (p<0.05), among which 48 proteins were overexpressed in V-24h and 59 proteins in E-24h. Together with distinctively expressed proteins shown in figure 2, a total of 90 proteins in V-24h and 138 proteins in E-24h were used for KEGG enrichment analysis (figure 5B). We matched 8 pathways in V-24h that are associated with the biosynthesis of amino acids, microbial metabolism in diverse environments, biosynthesis of secondary metabolites and carbon metabolism. Proteins involved in biosynthesis of amino acid are responsible for the production of branched-chain amino acids, threonine, arginine, glutamine, proline and tryptophan. In E-24h, only six pathways were enriched: mismatch repair, nucleotide excision repair, homologous recombination, DNA replication, pyrimidine metabolism and metabolic pathways. Again, this result demonstrates that DNA repair related pathways are main contributors to enrofloxacin-induced persisters.

**Figure 5.**
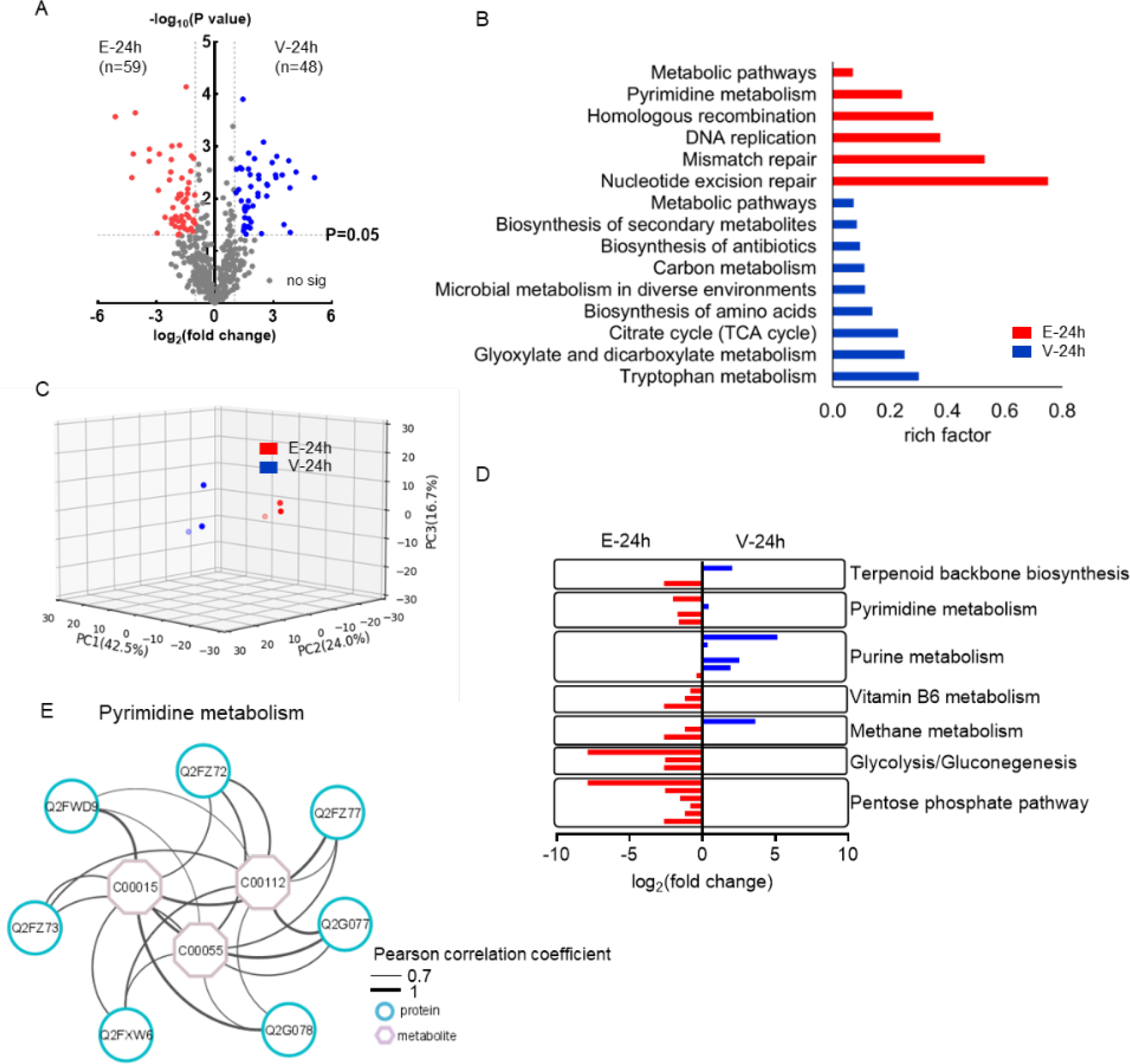
Proteins and metabolites comparison between persistent populations generated by vancomycin or enrofloxacin exposure. (**A**) T-test of shared proteins between V-24h and E-24h (n=483). (**B**) KEGG enrichment analysis was performed separately for V-24h and E-24h with significantly expressed proteins plus uniquely expressed proteins. Only pathways with adjusted p-value<0.05 were shown. (**C**) PCA of 95 identified metabolites present in V-24h (blue) and E-24h (red). The plot displays two distinct clusters, indicating good correlation between replicates and clear metabolic profile difference between V-24h and E-24h. (**D**) KEGG enrichment analysis with 95 metabolites (adjusted p-value <0.05). For comparison between V-24h and E-24h the average intensity was normalized with V-24h and shown as log_2_ fold change. Each bar represent one enriched metabolite. (**E**) As a common enriched pathway, the interaction map of proteins and metabolites involved in pyrimidine metabolism was analyzed. Only edges and nodes with pearson correlation cofficient ≥0.7 are shown. The line thickness represent the correlation value between 0.7 – 1.0. The blue circles represent proteins and pink octagons are metabolites.

Then, metabolomics was performed with 24h-treated persistent populations. Principal component analysis (PCA, figure 5C) shows qualified replicates and distinct profile in V-24h and E-24h samples. KEGG analysis resulted in eight significantly enriched pathways (adjusted P-value<0.05): metabolic pathways, pentose phosphate pathway, glycolysis/gluconeogenesis, methane metabolism, vitamin B6 metabolism, purine metabolism, pyrimidine metabolism and terpenoid back bone biosynthesis. The average intensity of each involved metabolite was normalized with V-24h and shown as log_2_ (fold change) (figure 5D), where positive values present higher intensity in V-24 (in blue) and negative values show more accumulation in E-24h (in red). Consistent with proteomics results that glycolytic enzymes were induced by vancomycin but repressed by enrofloxacin, vancomycin-induced persisters had relatively less accumulation of D-fructose-1,6-bisphosphate, glyceraldehyde-3-phosphate, and D-glycerate-2-phosphate. In purine metabolism, higher amounts of GMP, dGDP and GDP in V-24h were detected. This also supports the hypothesis that more ppGpp was generated, and by that, the stringent response was activated to combat vancomycin exposure. In E-24h the annotated metabolites in pyrimidine metabolism accumulated, which might favors DNA synthesis to respond against enrofloxacin exposure. As a common enriched pathway, the interaction network of pyrimidine related proteins and metabolites was depicted (figure 5E). Positive Pearson correlations were observed between the changes in the identified metabolites and proteins, indicating that these changes were likely interrelated.

### Differences in persistence mechanisms suggests new insights in how to eliminate persisters

Above we compared the molecular profiles of persistent populations induced by vancomycin or enrofloxacin, revealing both similarities and differences. As depicted in figure 6A, both vancomycin and enrofloxacin triggered persisters with more chaperones, elevated cell wall biosynthesis, less ribosomes and altered carbon metabolism. Meanwhile, vancomycin also stimulates the stringent response and potentially altered membrane composition by more generation of unsaturated fatty acid and branched-chain amino acids. For enrofloxacin persisters, DNA repair pathways and ABC transporters were more prominently observed.

**Figure 6.**
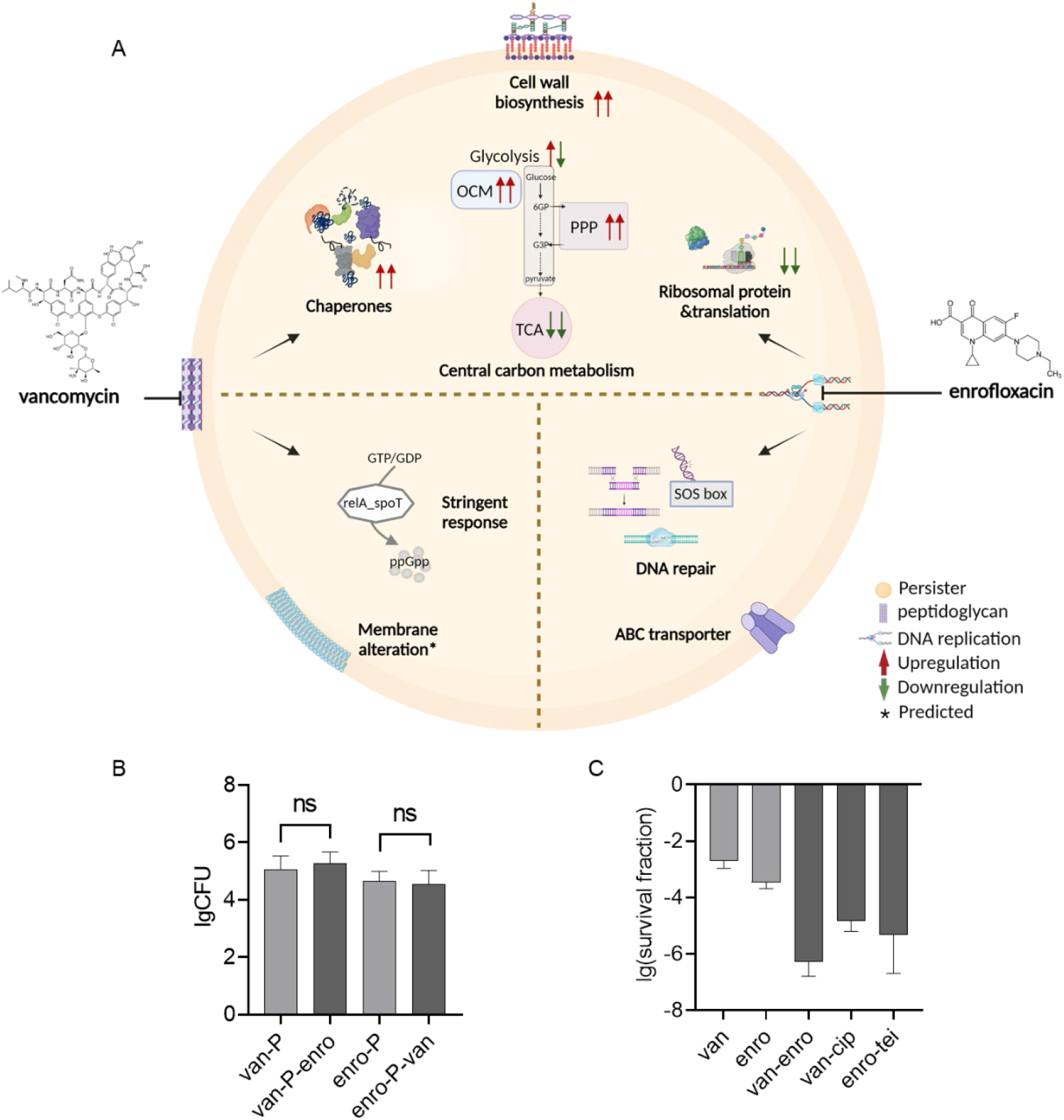
A. Scheme of the similar and unique mechanisms of persister formation induced by vancomycin and enrofloxacin. For upper area showing commonly altered pathway, arrows indicate the upregulation (red) or downregulation (green) of relevant pathways in either vancomycin persisters (left arrow) or enrofloxacin persisters (right arrow). The lower parts indicate unique pathways that are involved in either vancomycin persisters (left) or enrofloxacin persisters (right). **B**. Cross-tolerance testing of *S. aureus* persisters reveals that the persistent population generated from a single drug remained unaffected after an additional two hours of cross-treatment with another drug. n=9. **C**. Treatment with a combination of drugs (50-fold MIC+50-fold MIC) resulted in more killing than the single treatment (100-fold MIC). The minimum detectable survival fraction was 10^-8^. OCM: one carbon metabolism. van: vancomycin; enro: enrofloxacin; cip: ciprofloxacin; tei: teicoplanin. n≥3.

By understanding the mechanisms that underlie persister formation and maintenance, we may gain insights into how to effectively eliminate these cells. We hypothesized that similarities in molecular signalling pathways related to persister growth arrest may lead to cross-tolerance. This was confirmed experimentally as exposure to enrofloxacin of vancomycine surviving cells and visa versa failed to kill more cells after the initial persister formation (figure 6B). Besides, the differences between persistent populations make combined therapy a potent strategy to avoid high level of persistence. As shown in figure 6C, combining vancomycin and enrofloxacin in one treatment resulted in a significant increase in cell death, over 1000 times more than each of the single treatments. This suggests that the pathways bacteria used to generate persisters in single treatment were either inefficient or disrupted by the addition of the other drug, e.g., DNA damage caused by enrofloxacin could potentially interfere with relevant gene expression which the bacteria apply to persist under vancomycin pressure. We then also demonstrated a similar synergistic effect for the combination of vancomycin and another fluoroquinolone antibiotic ciprofloxacin, as well as with enrofloxacin combined with cell wall targeting teicoplanin. This corrobrates the potential of combination therapy to effectively reduce persisters rather than relying on sequential use of different antibiotics.

## Discussion

In this work, we collected samples during the biphasic kill curve to reveal mechanisms associated with persister formation and maintenance. Only 0.1% cells of initial cultures become persisters, as shown in the kill curve. Because of the eradication of the non-persisters after 24 hours of either vancomycin or enrofloxacin exposure, the remaining population consisted or 75% persisters, as was shown by flow cytometry Thus, the persisters were the major population of cells that we studied. Our hypothesis is that 24 hours incubation led to the lysis of most dead cells and the debris was washed out during sample preparation. Therefore, it is reasonable to say that proteins and metabolites in V-24h and E-24h mainly came from persistent cells.

Persisters, for decades, had been considered as subpopulations of bacteria without measurable metabolic activity. However, increased research evidenced that while persistent populations are non-dividing, certain metabolic pathways are still active to combat stress and to retain viability waiting for environmental conditions to improve, and resuscitation to take place. Our results also demonstrate that persisters are, to a certain level, metabolically active. Even under long-term stress from high levels of bactericidal antibiotics, persisters are still able to consume glucose, synthesize new proteins and regulate several metabolic pathways. In enrofloxacin triggered persisters, especially, we identified the unique expression of ABC transporters. This suggests that active extrusion of enrofloxacin may also plays a role in generating persistence and likely benefits the cells afterwards resuscitation as well.

Additionally, metabolic activity levels in two persistent populations induced by different antibiotics are likely different. Straightforwardly, as shown in figure 3C, vancomycin persisters are more active in glucose consumption and acetate production than enrofloxacin persisters. This hypothesis can also be inferred from the level of chaperones that aid in preventing protein aggregates, and ribosomes. A correlation between protein aggregation level and cell dormancy was proposed [53]. Specifically, controlled protein aggregation is suggested to facilitate persister formation by inactivating unnecessary metabolic activities, but excessive aggregation leads to deeper dormancy and even cell death. Thus, the different chaperone expression and aggregated protein levels observed in samples treated with vancomycin and enrofloxacin suggest that greater efforts were needed to sustain a specific level of protein aggregation in vancomycin-treated samples, particularly at V24-h, which could be one of the contributors to stimulate active carbon metabolism. Another indicator is the varying alteration in ribosomal proteins expression between two persistent populations. Wood et. al [54] discussed that while repressing ribosomes contribute to persister formation by reducing protein translation to force cells to enter dormancy, a certain intracellular ribosome level is essential for persisters to survive and resuscitate. With proteomics, we found lower expression of ribosomal proteins in both treated populations compared with untreated cells, but V-24h contained higher levels of ribosomal protein than E-24h. Overall, vancomycin persisters exhibit potentially higher metabolic activity than enrofloxacin persisters, which could also cause a difference in resuscitation rate.

One interesting finding is that the vancomycin triggered alterations mostly happened within one hour of antimicrobial incubation while most changes in enrofloxacin treated samples appeared only after four hours of exposure. In terms of the modes of action, vancomycin directly targets the cell wall while enrofloxacin needs to invade cells before inhibiting DNA synthesis. While this does not necessarily determine the killing efficiency of vancomycin and enrofloxacin, the fast stress responses of bacteria could be due to the instantaneous interaction between vancomycin and the cell wall. It is worth to note that the sampling timepoint might shape the conclusion in terms of persister formation mechanisms and even the efficacy of persister eradication due to the distinct stress response ‘reaction time’ of certain pathways or specific enzymes. Time resolved analysis is thus highly advocated.

Furthermore, our omics data provided valuable insights into the biophysical characteristics of persisters that might contribute to persister formation. One notable finding is that during type II

fatty acid biosynthesis, the repression of FabI in V-24h that could lead to generation of more unsaturated fatty acid and result in a more fluid membrane [55,56]. Similarly, higher production of branched-chain fatty acid also helps to maintain membrane fluidity [57]. Moreover, perturbed cell wall biosynthesis may lead to thicker cell walls as well as changed cell size. The results also hints the increased risk of resistance development during resuscitation as persisters contains higher level of resistance accelerating VraR, Pbp2 and SOS response-related proteins [58–60]. Still, further validation studies, e.g., using specific mutants and pure persistent population, are required for more detailed causal mechanistic analysis. This study, identifying molecular physiology that adapt *S. aureus* to become persisters when exposed to two types of antibiotics, is highly clinically relevant for preventing antibiotic failure. Noticeably, in this study, two types of antibiotics that triggered similar persister levels led to distinctive proteome and metabolome profiles. These suggest that persisters at the molecular physiological level are highly stress specific, meaning molecular features found in a population of persisters induced by one stressor could be different, or even opposite from those induced by another. In this case, the efficiency of an anti-persister strategy active against persisters induced by one antibiotic will not be guaranteed effective against a persistent population generated by another. Therefore, combinations of antibiotics can be sought to mutually inactivate the mechanisms which the bacteria utilize to generate persisters against either of the antibiotics in single treatment. This way, persister formation may be prevented, allowing proper treatment of patients and eliminating the entire reservoir of the infection. Future studies should focus on more detailed mechanistic analyses, preferably in isolated persisters, focussing on the use of mutant strains to go from the current molecular characterisation and correlation to causative relations.

## Materials and Methods

### Strain cultivation and time-kill kinetics assay

*Staphylococcus aureus* ATCC 49230 strain (UAMS-1, a highly virulent clinical isolation from patient with chronic osteomyelitis) was used in this experiment and was incubated at 37 °Cwith shaking at 200 rpm, unless otherwise specified. Overnight culture from a single colony cultured in tryptic soy broth (TSB) was 1:100 inoculated into fresh TSB medium and incubated until early log phase (OD_600_=0.4-0.6). The culture was then diluted with fresh TSB to OD_600_=0.2 to make consistent starting amount. Diluted culture was divided into three Erlenmeyer flasks for three treatment groups. One of the cultures without antibiotic exposure was used as control. For the other two flasks, antibiotics were added separately to a final concentration of 100-fold MIC for single treatment or 50-fold MIC for combined treatment. MIC measurement and the quantification of CFU were performed following the previous protocol [61]. The MICs of each antibiotic (all purchased from Sigma-aldrich Chemie NV, the Netherland) are vancomycin 1 μg/ml, enrofloxacin 0.25 μg/ml, teicoplanin 0.25 μg/ml, ciprofloxacin 1 μg/ml.

For time-kill kinetics assay, aliquots of 1ml of culture were removed before and after 1h, 3h, 5h and 24h of antibiotic exposure, and washed twice with phosphate-buffered saline (PBS) buffer to remove antibiotics. Surviving cells in each sample were then quantified by quantitative culture of 10-fold serial dilutions with PBS plated on TSB agar plate. After overnight incubation, the number of colonies was quantified, and the surviving fraction of treated samples compared with control was calculated and visualized as kill curve. For cross tolerance, 100-fold MIC of enrofloxacin or vancomycin was added directly into 24 hours vancomycin or enrofloxacin treated samples, respectively. Then, two hours incubation was performed before quantification. To detect the proportion of persister cells after antibiotic exposure, CFDA-PI double staining and subsequent flow cytometry was used for control and 24h treated cultures (unpublished method).

### Extracellular metabolites measurement

Extracellular medium was filtered by 0.22 μm filters before and during 1h, 2h, 3h, 4h and 24h antibiotic incubation. The standards for the High-Performance Liquid Chromatography (HPLC) measurement of glucose and acetate were prepared with a set of concentration within 0-50 mM. All standards and samples were measured by HPLC (LC-20AT, Prominence, Shimadzu) equipped with Ion exclusion Rezex ROA-Organic Acid H+(8%) column (300x 7.8 mm; Phenomenex), along with a guard column (Phenomenex). The injection volume was set to 50 μl using the SIL-20AC autosampler (Shimadzu), and the system operated at a pressure of 50 bar. Aqueous H_2_SO_4_ (5 mM) was used as mobile phase at a flow rate of 0.5 ml/min at 55 °C. The UV detection was performed at λ=210 nm using the SPD-20A UV/VIS detector, and a refractive index detector was also utilized (RID 20A, Shimadzu).

### Samples collection

15 OD cultures of vancomycin or enrofloxacin treated groups were collected during incubation to isolate protein and metabolites. For proteins, samples were collected before and after 1h-, 4h-, 24h-antibiotic exposure. Metabolites from final 24h-treated samples were also extracted for metabolomics. The isolation protocol is modified from [62]. Specifically, after filtration and washing twice with ice-cold 0.6 % NaCl solution, filter paper was quickly removed into a 50 ml Falcon tube containing 3 ml ice-cold 60 % ethanol as extraction solution. After vigorously vortexing for about ten seconds, the tube was dropped into liquid nitrogen to quench the metabolism. Samples were stored in -80 °C until all triplicate treatment groups were collected and ready for protein and metabolites isolation.

Frozen samples were thawed on ice and mixed by vortexing for ten second to detach the cells from the filter. Afterwards, 1ml of the resulting cell suspension was transferred into a 2 ml homogenizer tube containing 0.5 ml 0.5 mm glass beads and each sample was aliquoted into three tubes. Cells were then disrupted by Bead Ruptor Elite Homogenizer (OMNI international, Georgia, America). During the beads beating, the ambient temperature was kept at around 0 °C. Next, cell suspensions from the same sample were transferred to a 15 ml tube. The glass beads were washed twice with ice-cold ethanol and the washing solution was also transferred to the same tube. To separate metabolites and proteins, tubes were centrifuged for 5 mins at 10,000 ⨯g. 4 ml supernatant containing metabolites was transferred and dried by a nitrogen stream. Nonpolar and polar metabolites were suspended by isopropanol: acetonitrile: water=4:3:1 and acetonitrile: water=1:1, respectively. The rest of supernatant and pellet with proteins was dried in a centrifugal vacuum evaporator and resuspend with 1 % SDS (sodium dodecyl sulfate, Duchefa Biochemie B.V. Haarlem, the Netherlands) in 100 mM ammonium bicarbonate (ABC, Sigma-aldrich Chemie NV, the Netherlands).

### LC-MS sample preparation and analysis

#### Metabolomics

The dry apolar metabolites obtained by extractions by monophasic methods were reconstituted in 200 μL water the dry polar metabolites were in 200 μL ACN/Water (1:1). 10-μl was injected for non-targeted metabolomics onto a CSH-C18 column (100 mm × 2.1 mm,1.7 μm particle size, Waters, Massachusetts, USA) for apolar metabolites analysis or a BEH-Amide column (100 mm × 2.1 mm,1.7 μm particle size, Waters, Massachusetts, USA) for a-polar metabolites by an Ultimate 3000 UHPLC system (Thermo Scientific, Dreieich Germany). Using a binary solvent system (A: 0.1% FA in water, B: 0.1% FA in ACN) metabolites were separated on the CSH-C18 column by applying a linear gradient from 1% to 99% B in 18 minutes at a flow rate of 0.4 ml/min, while polar metabolites were separated on the BEH-Amide column by applying a linear gradient from 99% B to 40% B in 6 minutes and then to 4% B in 2 minutes at a flow rate of 0.4 ml/min. Eluting analytes were electrosprayed into a hybrid trapped-ion-mobility-spectroscopy quadrupole time of flight mass spectrometer (tims TOF Pro, Bruker, Bremen Germany), using a capillary voltage of 4500 volt in positive mode and 3500 volt in negative mode, with source settings as follows: end plate offset 500 volt, dry temp 250 °C, dry gas 8 l/min and nebulizer set at 3 bar both using nitrogen gas. Mass spectra were recorded using a data dependent acquisition approach in the range from m/z 20-1300 for polar and 100-1350 for the apolar metabolites in positive and negative ion mode using nitrogen as collision gas. Auto MS/MS settings were as follows: Quadupole Ion Energy 5 eV, Quadrupole Low mass 60 m/z, Collision Energy 7 eV. Active exclusion was enabled for 0.2 min, reconsidering precursors if ratio current/previous intensity > 2.

#### Proteomics

The pellets obtained above were dissolved in 300μl 1% SDS in 100mM ABC and vortexed thoroughly. The BCA assay was used to determine the concentration of protein according to the manual and TCEP and CAA added to 10 mM and 30 mM, respectively, and the mix incubated for 0.5 h at room temperature. Samples were processed using the SP3 protein clean up [61] and trypsin (protease/protein, 1:50, w/w) was added and protein was digested at 37°C overnight. The supernatant was acidified with formic acid (FA) (1% final concentration and a pH ∼2), and while on a magnetic stand to trap magnetic beads the supernatant was moved to a clean tube. For LC-MS analysis ∼ 200ng (measured by a NanoDrop at a wavelength of 215 nm) of the peptide was injected by an Ultimate 3000 RSLCnano UHPLC system (Thermo Scientific, Germering, Germany).

Following injection, the peptides were loaded onto a 75um x 250 mm analytical column (C18, 1.6 μm particle size, Aurora, Ionopticks, Australia) kept at 50°C and flow rate of 400 nl/min at 3% solvent B for 1 min (solvent A: 0.1% FA, solvent B: 0.1% FA in ACN). Subsequently, a stepwise gradient of 2% solvent B at 5 min, followed by 17% solvent B at 24 min, 25% solvent B at 29 min, 34% solvent B at 42 min, 99% solvent B at 33 min held until 40 min returning to initial conditions at 40.1 min equilibrating until 58 min. Eluting peptides were sprayed by the emitter coupled to the column into a captive spray source (Bruker, Bremen Germany) which was coupled to a TIMS-TOF Pro mass spectrometer. The TIMS-TOF was operated in PASEF mode of acquisition for standard proteomics. In PASEF mode, the quad isolation width was 2 Th at 700 m/z and 3 Th at 800 m/z, and the values for collision energy were set from 20-59 eV over the TIMS scan range. Precursor ions in an m/z range between 100 and 1700 with a TIMS range of 0.6 and 1.6 Vs/cm2 were selected for fragmentation. 10 PASEF MS/MS scans were triggered with a total cycle time of 1.16 seconds, with target intensity 2e4 and intensity threshold of 2.5e3 and a charge state range of 0-5. Active exclusion was enabled for 0.4 min, reconsidering precursors if ratio current/previous intensity >4.

## Data analysis

The metabolites mass spectrometry raw files were submitted to MetaboScape 5.0 (Bruker Daltonics, Germany) used to perform data deconvolution, peak-picking, and alignment of m/z features using the TReX 3D peak extraction and alignment algorithm (EIC correlation set at 0.8). All spectra were recalibrated on an internal lockmass segment (NaFormate clusters) and peaks were extracted with a minimum peak length of 7 spectra (8 for recursive extraction) and an intensity threshold of 500 counts for peak detection. In negative mode ion deconvolution setting, [M-H]-was set for primary ion, seed ions were [M+Cl]- and common ions [M-H-H2O]-, [M+COOH]-. For positive mode, the primary ion was [M+H] +, seed ions were [M+Na] +, [M+K] +, [M+NH4] + and [M-H-H2O] + were common ions. Features were annotated, using SMARTFORMULA (narrow threshold, 3.0 mDa, mSigma:15; wide threshold 5.0 mDa, mSigma:30), to calculate a molecular formula. Spectral libraries including Bruker MetaboBASE 3.0, Bruker HDBM 2.0, MetaboBASE 2.0 in silico, MSDIAL LipidDBs, MoNA VF NPL QTOF, AND GNPS export, were used for feature annotation (narrow threshold, 2.0 mDa, mSigma 10, msms score 900, wide threshold 5.0 mDa, mSigma:20 msms score 800). An annotated feature was considered to be of high confidence if more than two green boxes were present in the Annotation Quality column of the program and low confidence if less than two green boxes were present.

After searching and peak-picking, positive and negative profiles were combined for further analysis. Metabolomic data analysis was carried out by using MetaboAnalyst 5.0 (https://www.metaboanalyst.ca/MetaboAnalyst/home.xhtml). At first, the data showing a poor variation were filtered on Inter Quartile Range (IQR) and then features were normalized by median normalization, scaled by Auto scaling and transformed to a logarithmic scale (base of 2).

Generated Mass spectra for pellets were analyzed with Maxquant (Version 1.6.14) for feature detection and protein identification. Tims-DDA was set in Type of Group specific and other parameters were set as the default. Search included variable modifications of methionine oxidation, and a fixed modification of cysteine carbamidomethyl and the proteolytic enzyme was trypsin with maximum 2 missed cleavage. A *S. aureus* database (version 2019 downloaded from Uniprot) was used for database searches. To improve mass accuracy of matching precursors, the “match between runs” option was applied within a match window time of 0.2 min and a match ion mobility window of 0.05. Proteins of label free quantification (LFQ) calculated for each pellet represented normalized peptide intensities correlating with protein abundances. Finally, all the quantification and annotation information were summed in in the output proteinGroup.txt.

Proteins and metabolites that were found in at least two replicates were considered to be reproducibly detected and used for analysis. Then data were analyzed by Perseus [63] for log_2_ transform, Pearson correlation coefficient analysis, hierarchical cluster analysis, PCA assay, Z-score normalization and statistical analysis. Significantly expressed proteins are those with |log_2_(fold change) |>1 and p value <0.05. The proteomic pathway analysis was done with KOBAS [64] and metabolic pathway enrichment was conducted on website MBROLE 2.0 (Metabolites Biological Role) [65], only pathways with adjusted p-value <0.05 were considered to be significant.

## Supporting information

Supplemental data insoluble proteins

All proteomics and metabolomics data as deposited in the public domain

## Data availability

All mass spectral data related to this publication have been deposited in the MassIVE repository (https://massive.ucsd.edu/ProteoSAFe/static/massive.jsp) under the dataset identifier MSVXXXX

## Author Contributions

Conceptualization, S.B., S.A.J.Z. and S.L.; performing experiments and analyzing data, S.L., Y.H., S.J., P.L., G.K.; writing—original draft preparation, S.L.; writing—review and editing, G.K. S.B. and

S.A.J.Z. All authors have read and agreed to the published version of the manuscript.

