## Supplemental data insoluble proteins for "Molecular physiological characterization of the dynamics of persister formation in *Staphylococcus aureus*"

The relationship between protein aggregation and antibiotic persistence has been recently discovered where aggregated protein contributes to antibiotic persistence possibly via inhibiting growth and other unnecessary metabolic activities, meanwhile, disaggregation by chaperon proteins is required for resuscitation. To detect protein aggregation level in triggered persisters, insoluble proteins were extracted from OD15 of samples before and at 1, 4 and 24h of enrofloxacin or vancomycin exposure. The volumes of loaded samples were normalized based on the total amount of protein in each sample. After sodium dodecyl-sulfate polyacrylamide gel electrophoresis (SDS-PAGE) and Coomassie blue staining, the result (figure S1) shows that both vancomycin and enrofloxacin exposure led to protein aggregation and vancomycin persisters contains higher level of aggregated protein compared with enrofloxacin persisters. Interestingly, vancomycin triggered gradually increased protein aggregation levels along the incubation but enrofloxacin led to slightly decrease of protein aggregation over time. Additionally, unique bands were visible only in the vancomycin-treated samples, indicating a distinctive protein profile in those cells.

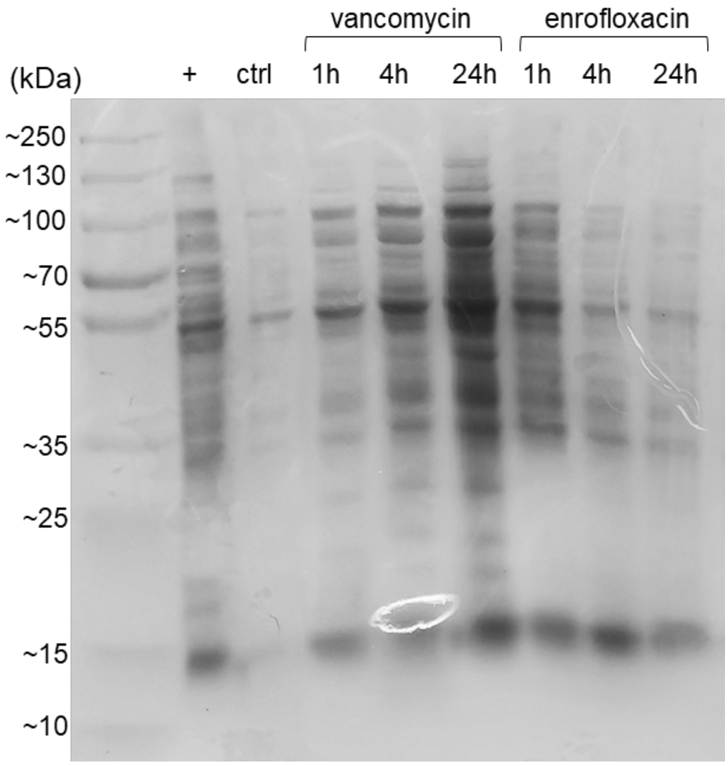


**Figure S1**. SDS-PAGE of insoluble proteins obtained from 42 ℃ incubated culture as positive control (ctrl+), untreated control samples (ctrl), and samples treated for 1, 4, and 24h by vancomycin and enrofloxacin. The loading volume of each sample was normalized based on the concentration of total protein.

### Method:

To visualize protein aggregation level, 15 OD of each sample were collected. The concentration of total protein of each sample was firstly quantified by beads beating with samples in 1 % SDS solution and Bicinchoninic acid assay for normalization. Insoluble protein was extracted and visualized following the protocol of Tomoyasu *et al*. [1]. Untreated cultures were used as control and overnight incubated cultures in 42 ℃ as positive control [2].
